# Noise induces intercellular Ca^2+^ signaling waves and the unfolded protein response in the hearing cochlea

**DOI:** 10.1101/2024.06.05.597671

**Authors:** Yesai Park, Jiang Li, Noura Ismail Mohamad, Ian R. Matthews, Peu Santra, Elliott H. Sherr, Dylan K. Chan

## Abstract

Exposure to loud noise is a common cause of acquired hearing loss. Disruption of subcellular calcium homeostasis and downstream stress pathways in the endoplasmic reticulum and mitochondria, including the unfolded protein response, have been implicated in the pathophysiology of noise-induced hearing loss. However, studies on the association between calcium homeostasis and stress pathways has been limited due to limited ability to measure calcium dynamics in mature-hearing, noise-exposed mice. We used a genetically encoded calcium indicator mouse model in which GcAMP is expressed specifically in hair cells or supporting cells under control of Myo15Cre or Sox2Cre, respectively. We performed live calcium imaging and UPR gene expression analysis in 8-week-old mice exposed to levels of noise that cause cochlear synaptopathy (98 db SPL) or permanent hearing loss (106 dB SPL). UPR activation occurred immediately after noise exposure and was noise dose-dependent, with the pro-apoptotic pathway upregulated only after 106 dB noise exposure. Spontaneous calcium transients in hair cells and intercellular calcium waves in supporting cells, which are present in neonatal cochleae, were quiescent in mature-hearing cochleae, but re-activated upon noise exposure. 106 dB noise exposure was associated with more persistent and expansive ICS wave activity. These findings demonstrate a strong and dose-dependent association between noise exposure, UPR activation, and changes in calcium homeostasis in hair cells and supporting cells, suggesting that targeting these pathways may be effective to develop treatments for noise-induced hearing loss.

## Introduction

Noise-induced hearing loss (NIHL) affects an estimated 40 million individuals in the US, with no approved medical treatments (1). A number of cellular mechanisms have been proposed to be involved in NIHL, including oxidative stress (2,3), JNK/ERK pathway activation (4), mitochondrial stress (5), endoplasmic reticulum (ER) stress, and the Unfolded Protein Response (UPR) (6); targeting these pathways have identified multiple candidate drugs that prevent NIHL to varying degrees in mouse: Ru360, a mitochondrial Ca^2+^ uniporter (MCU) inhibitor that reduces mitochondrial Ca^2+^ uptake and overload (7); ISRIB, an eIF2B activator that inhibits the pro-apoptotic PERK/CHOP pathway of the UPR (6); D-JNKI-1, a peptide inhibitor of c-Jun N-Terminal Kinase that blocks the MAPK-JNK signaling pathway (8); and N-acetyl cysteine, an antioxidant that reduces reactive oxygen species and oxidative stress (9). Of these, D-JNKI-1 and NAC have been tested in humans and shown limited efficacy in reducing NIHL (10,11). Better understanding of the precise mechanistic pathways by which noise induces cellular pathways in the cochlea leading to hair-cell death is essential to develop more effective treatments.

Many of these potential pathways – mitochondrial/oxidative stress, and ER stress and the UPR — are activated upon disruption of Ca^2+^ homeostasis. In addition to acquired hearing loss, many genetic forms of deafness involve molecules involved in Ca^2+^ flow and homeostasis in cochlear cells (12), illustrating the broad-based importance of these pathophysiologic mechanisms. Dysregulation of subcompartmental Ca^2+^ homeostasis has been directly implicated in hair-cell death using an aminoglycoside model of ototoxicity in zebrafish hair cells, in which ER Ca^2+^ depletion leads to cytosolic Ca^2+^ accumulation, mitochondrial Ca^2+^ overload, and mitochondrial stress (13,14). In mammals, both ER Ca^2+^ depletion (leading to UPR activation) and mitochondrial Ca^2+^ overload (leading to oxidative stress) have been implicated in hearing loss (6, 8). We have shown that disruption of TMTC4, an ER-resident, hair-cell-specific gene implicated in progressive hearing loss in mice and humans (15), causes ER Ca^2+^ depletion and UPR activation, and that noise exposure causes UPR activation (6). Importantly, we found that targeting the UPR with ISRIB, a small molecule activator of eIF2B, reduces noise-induced hearing loss and cochlear synaptopathy (6,16). On the other hand, genetic or pharmacologic disruption of MCU, which reduces mitochondrial Ca^2+^ accumulation, also protects against NIHL (7). Despite this strong evidence that disruption of Ca^2+^ homeostasis in hair cells can lead to hair cell death through ER- and/or mitochondrial stress pathways, it is not clear how, and whether, noise trauma directly causes this Ca^2+^ dysregulation in hair cells.

One possibility is that mechanical trauma or noise exposure affects Ca^2+^ homeostasis through induction of intercellular Ca^2+^ signaling (ICS) waves in supporting cells. ICS waves have been studied extensively in the central nervous system; they propagate across glial networks, where they respond to mechanical and excitotoxic trauma and mediate neuronal repair, death, and migration (17,18). Noise and mechanical trauma have also been suggested to induce ICS waves across supporting-cell networks in the cochlea (19), but not hair cells; ICS waves are dependent on Cx26 to propagate, and Cx26 is only expressed in supporting cells (20). These changes in Ca^2+^ flux have been implicated in downstream cellular signaling cascades, including activation of the ERK pathway (21) as well as the UPR (6). In the neonatal cochlea, ICS waves occur spontaneously in supporting cells of the inner sulcus and outer sulcus in the late stages of cochlear development, synchronizing inner-hair-cell firing (22,23). ICS waves can also be triggered in neonatal cochleae by external ATP or direct mechanical trauma (19,26). Though spontaneous ICS activity was initially thought to become quiescent after the onset of hearing (23), subsequent studies have shown limited evidence of spontaneous (24) and noise-evoked (CR16) ICS waves in the adult mouse and gerbil cochlea, respectively. The role of these supporting-cell ICS waves in the inner ear’s response to noise, however, is poorly understood.

Despite these studies demonstrating links between noise exposure and ICS signaling in supporting cells in mechanically traumatized neonatal cochlea (19), between Ca^2+^ transients in hair cells and cell death in zebrafish aminoglycoside ototoxicity (13,14), and between ER and mitochondrial stress and noise exposure in mice (6,7), comprehensive evaluation of the Ca^2+^ and stress pathways by which noise exposure leads to hearing loss has been limited due to the absence of a single experimental model in which dynamic cellular processes can be observed live in the mature, hearing cochlea after physiologically relevant noise exposure. Such work would require a single model system in which the effect of noise exposures on Ca^2+^ homeostasis in hair cells and supporting cells can be directly correlated with the effect of the same noise exposures on downstream pre-apoptotic pathways, such as the UPR, and hearing loss.

In this study, we sought to evaluate the hypothesis that excessive noise induces Ca^2+^ homeostatic changes in both hair cells and supporting cells, leading ultimately to pro-apoptotic UPR activation. In order to do this, we have developed a live-imaging model of the mature, hearing cochlea, enabling direct visualization of cytosolic Ca^2+^ dynamics in hair cells and supporting cells after physiologically-relevant noise exposure. This model allows us to measure three elements of the Ca^2+^ dysregulation/UPR axis in response to the same set of physiologically-relevant noise exposures that cause either temporary or permanent shifts in hearing thresholds: 1) UPR gene expression; 2) Ca^2+^ transients in hair cells; and 3) ICS waves in supporting cells. Evidence of these three phenomena provides support that they are all connected in the early response of the cochlea to acoustic overstimulation.

## Results

### UPR expression after NIHL

We first investigated the response of the UPR to varying levels of noise exposure. 8-week-old male and female wild-type CBA/J mice were exposed to 8-16 kHz octave-band noise for 2h at three levels: 94 dB SPL, which does not cause any threshold shift; 98 dB SPL, which causes temporary threshold shift (TTS) and cochlear synaptopathy (16); and 106 dB SPL, which causes permanent threshold shift (PTS) and hair-cell death (6) (**Supplementary Figure S1**). Cochleae were extracted from mice 2 hours after completion of noise exposure, and expression levels of three UPR marker genes – BiP, CHOP, and S-XBP1 – were measured by qPCR (**Figure 1**). BiP, a marker for general activation of the UPR, was significantly elevated with all noise exposure levels, demonstrating that noise exposure upregulates the UPR. No significant changes in S-XBP1, a specific marker for the pro-homeostatic arm of the UPR, were seen compared to control. CHOP, a marker of the pro-apoptotic arm of the UPR, was elevated only in male mice exposed to 106 dB SPL. However, when the ratio of CHOP/S-XBP1, which indicates a shift in the balance of the UPR towards apoptosis, was compared across conditions, significant elevation of this ratio was seen in both male and female mice exposed to 106 dB SPL noise. These results demonstrate that although the UPR is activated overall with any noise exposure (as indicated by BiP), noise levels that cause PTS and hair-cell loss are associated specifically with significant elevation of the CHOP/S-XBP1 ratio. Overall, no statistically significant differences were seen relating to biological sex; for this reason, all subsequent experiments were pooled across sexes.

**Figure 1.**
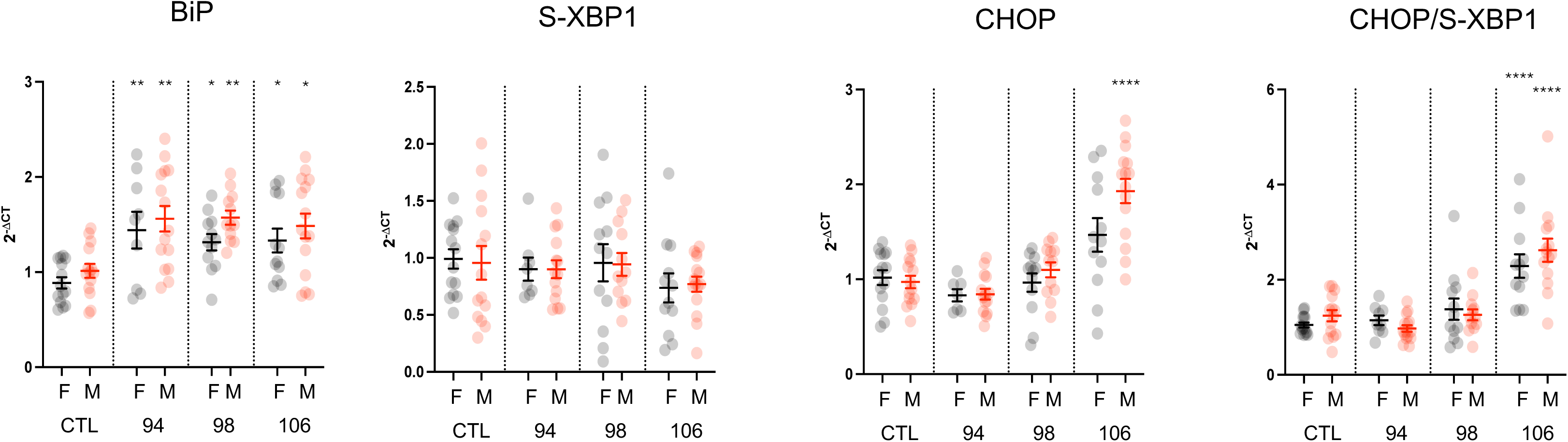
Noise exposure induces UPR upregulation in the cochlea *in vivo*. To investigate the sound level dose-dependence of UPR activation after acoustic overstimulation, we exposed 8-week-old male and female wild-type CBA/J mice to 8-16 kHz octave-band noise for 2 hours at 94, 98 106, levels that respectively induce no hearing loss, temporary threshold shift (TTS) with cochlear synaptopathy, and permanent threshold shift (PTS) with hair-cell death, respectively. Cochlea were harvested for qPCR measurement of BiP, S-XBP1, and CHOP mRNA expression using the 2^-DCT^ method relative to GAPDH expression and normalized to control (non-noise-exposed) levels. Data were cleaned by removing outliers (ROUT method, Q=1%) and compared with one-way ANOVA with Dunnett’s test for multiple comparisons against control for each condition. * p < 0.05; ** p < 0.01; *** p < 0.001; **** p < 0.0001. N = 14 (CTL-F); 14 (CTL-M); 9 (94 dB – F); 15 (94 dB – M); 12 (98 dB – F); 12 (98 dB – M); 12 (106 dB – F); 15 (106 dB – M).

We next investigated the time-course of UPR activation after 98 and 106 dB SPL noise exposure, our two primary models for TTS and PTS, respectively. Cochleae were harvested at 0, 2, 12, and 24h after completion of the 2h noise exposure, as well as 2 weeks later, and compared with control, non-noise-exposed animals (**Figure 2**). Elevation in BiP was seen immediately after noise exposure, followed by changes in CHOP and S-XBP1. Elevation in CHOP/S-XBP1 ratio peaked at 2h after noise exposure and was more pronounced in mice exposed to the louder 106 dB SPL noise. These findings demonstrate that UPR gene expression changes are an early and dose-dependent response to noise exposure in mice.

**Figure 2.**
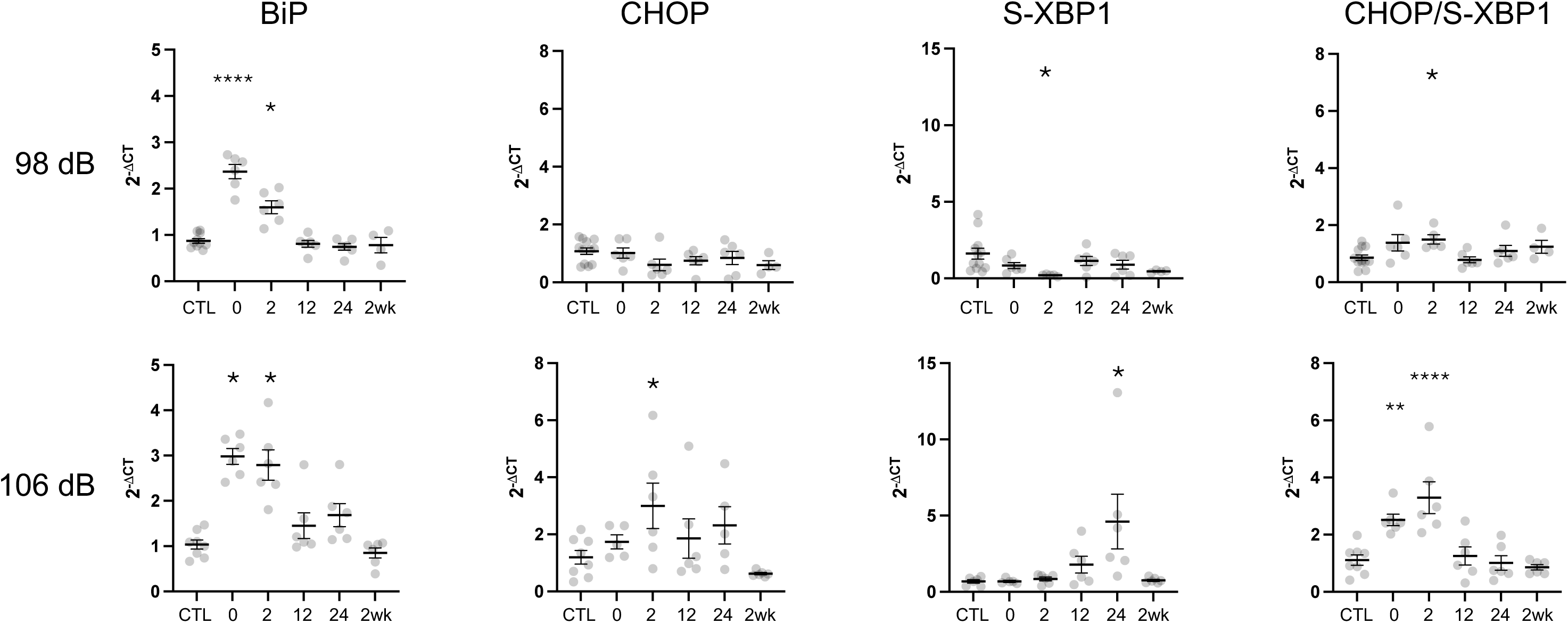
Noise exposure induces rapid UPR upregulation in the cochlea *in vivo.* To investigate the temporal evolution of the UPR after acoustic overstimulation, we exposed 8-week-old male wild-type CBA/J mice to 8-16 kHz octave-band noise for 2 hours at 98 dB SPL (top), which induces TTS, or 106 dB SPL (bottom), which induces PTS. Cochleae were harvested at the indicated timepoints after noise exposure for qPCR measurement of BiP, S-XBP1, and CHOP mRNA expression using the 2^-DCT^ method relative to GAPDH expression and normalized against control (non-noise-exposed) levels. Data were cleaned by removing outliers (ROUT method, Q=1%) and compared with one-way ANOVA with Dunnett’s test for multiple comparisons against control for each condition. * p < 0.05; ** p < 0.01; *** p < 0.001; **** p < 0.0001. N = 12 (CTL – 98 dB); 8 (CTL – 106 dB); 6 (0, 2, 12, 24h, 98 dB and 106 dB); 6 (2wk – 106 dB); 4 (2wk – 98 dB).

### Live Ca^2+^ imaging in hair cells and supporting cells in the neonatal cochlea

We then sought to use these same noise-exposure models —98-dB and 106-dB, 8-16 kHz octave-band noise, which are associated with TTS (and UPR activation without shift towards apoptosis) and PTS (with pro-apoptotic UPR activation), respectively — and directly visualize Ca^2+^ dynamics in the organ of Corti. However, live Ca^2+^ imaging had previously been performed primarily in neonatal cochlear cultures, which cannot be stimulated with sound, and prior instances of Ca^2+^ imaging in the adult cochlea (24,26) used exogenous dyes that did not sufficiently label hair cells. We therefore developed an acute explant preparation of the temporal bone from juvenile, mature-hearing mice (27) expressing the cytosolic Ca^2+^ indicator GcAMP6s in a Cre-dependent manner in supporting cells (driven by Sox2Cre) or hair cells (driven by Myo15Cre). This preparation was similar to one previously described that demonstrated ICS waves in the adult mouse using exogenous Fluo-4-AM dye for Ca^2+^ sensing (24), but instead uses a genetically-encoded Ca^2+^ indicator to further expedite imaging after euthanasia and enable both hair-cell and supporting-cell-specific labeling.

To validate this model, we first confirmed expression of GcAMP and validated its ability to detect Ca^2+^ activity in supporting cells and hair cells in neonatal mice. Neonatal cochlear cultures from Sox2Cre-GcAMP mice expressed GcAMP in supporting cells, but not hair cells (**Figure 3**), and exhibited spontaneous ICS waves identical to those seen using exogenous fluorophores (FURA-2 (19) and Fluo-4 (22,24)). ICS waves were observed in Kölliker’s organ at the inner sulcus (IS), as well as in outer sulcus (OS) (**Supplemental Video 1, Figure 3**). Overall, ICS waves propagated at 15.5 ± 0.5 μm/s (mean ± sem from 128 waves in 15 cochleae) in the IS and 27.9 ± 0.4 μm/s (N=914 waves) in the OS, and occurred at a rate of 0.03 waves/s (IS) and 0.20 waves/s (OS). Drugs that prevent cytosolic Ca^2+^ clearance — vanadate, which blocks extrusion through PMCA, and thapsigargin, which blocks ER re-uptake through SERCA — affected both single-cell-level Ca^2+^ peaks and ICS characteristics. Vanadate, but not thapsigargin, increased the frequency of Ca^2+^ peaks and ICS waves (**Figure 3B,E**). Both drugs significantly increased the decay time for the cytosolic Ca^2+^ peak to return back to baseline (**Figure 3C-D**) as well as the distance of ICS wave propagation (**Figure 3F**). Finally, both drugs also increased steady-state cytosolic Ca^2+^ levels (**Figure 3G-H**). Taken together, these findings suggest that supporting-cell-specific GcAMP signal in the Sox2Cre-GcAMP model is accurately representing ICS wave activity.

**Figure 3.**
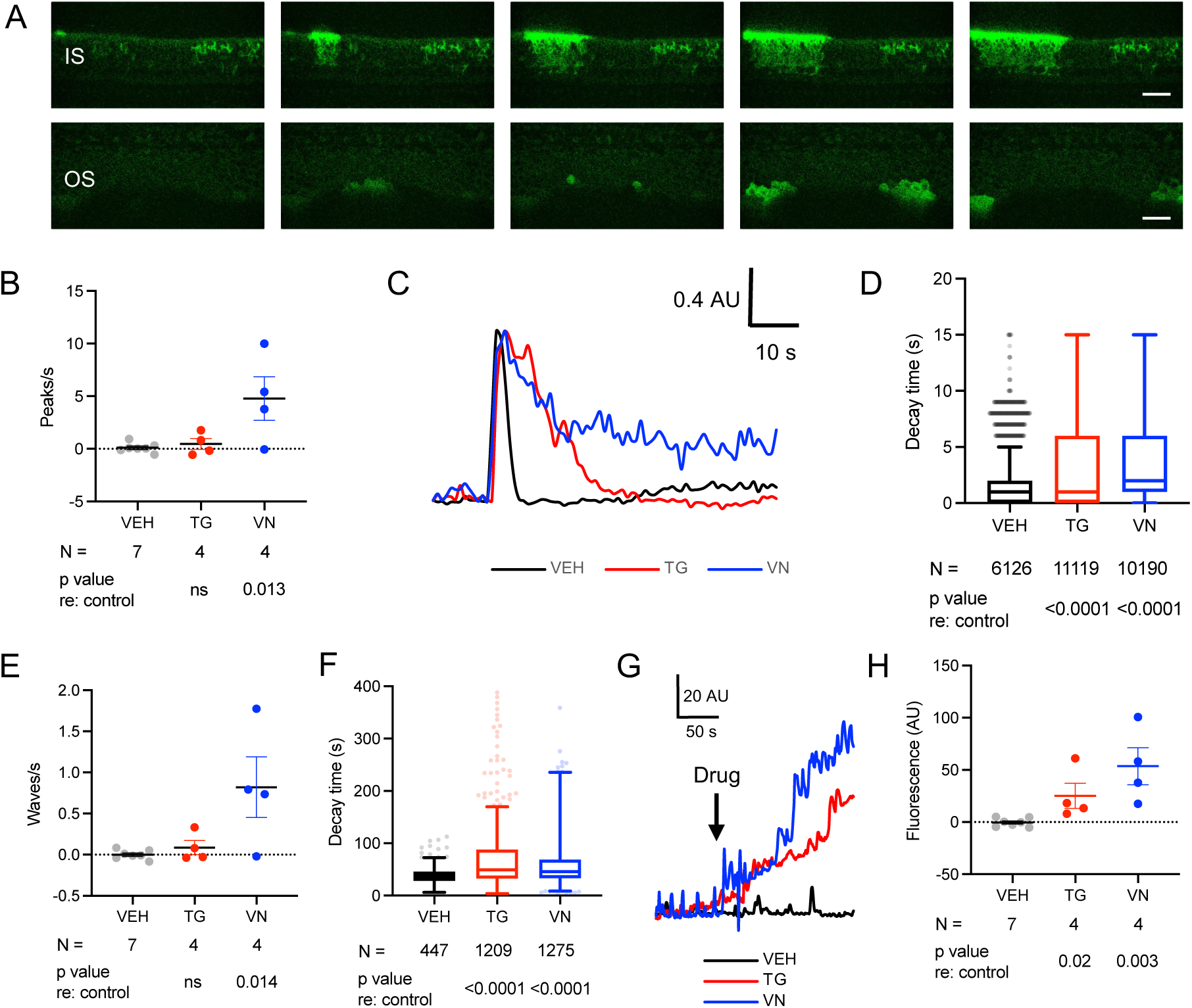
Ca^2+^ activity in neonatal cochlear supporting cells. **A. Live imaging of Sox2Cre-GcAMP neonatal cochlea.** In supporting cells of the inner sulcus (IS) and outer sulcus (OS) of the neonatal cochlea, spontaneous intercellular Ca^2+^ waves are observed. Time interval between successive images is 1s. Scale bar: 20 μm. **B. Change in number of Ca^2+^ peaks per second after drug treatment.** Compared to baseline (media only), no change in frequency of Ca^2+^ peaks is seen with vehicle (VEH, black) or thapsigargin (TG, red); vanadate (VN, blue) induced increased Ca^2+^ peak activity. **C-D. Ca^2+^ decay time with drug treatment.** Fluorescence levels **(C)** and peak decay time **(D)** demonstrate significant prolongation of return to baseline Ca^2+^ levels in the presence of TG or VN. In **(C),** peak height is normalized to maximum amplitude for each peak. **E. Change in number of ICS waves per second after drug treatment.** No change in frequency of ICS waves is seen with vehicle (VEH) or thapsigargin (TG); vanadate (VN) induced more ICS activity. **F. ICS wave distance propagation with drug treatment.** ICS waves travelled significantly farther in the presence of TG, but not VN. **G-H. Change in steady-state Ca^2+^ after drug treatment.** Fluorescence levels **(G)** and mean amplitude **(H)** over a 300s recording period, representing steady-state Ca^2+^ level across the entire cochlea, increased after TG and VN, but not VEH application. **B, E, H:** Means ± sem, with individual cochlea-level values as dots. **D and F:** Tukey plots representing all peaks **(D)** or waves **(F)** measured under the indicated conditions. P values are as indicated for pairwise comparisons versus control (VEH) on 2-tailed unpaired Student’s t test. ns, not significant. AU, arbitrary units.

We then examined neonatal cultures from Myo15Cre-GcAMP mice, which express GcAMP in hair cells with scant off-target labelling (**Supplemental Video 2**, **Figure 4).** Minimal spontaneous activity was observed (**Supplemental Video 2**); occasional spontaneous Ca^2+^ transients were observed in IHCs, but these never propagated as ICS waves, consistent with the fact that HCs do not express connexin 26 necessary for the direct and paracrine signaling that underlies wave propagation (20). Application of ATP, however, induced a large cytosolic Ca^2+^ transient in all hair cells, demonstrating an intact purinergic response; clearance of these ATP-induced Ca^2+^ peaks, as well as steady-state cytosolic Ca^2+^ level, was sensitive to vanadate and thapsigargin (**Figure 4B-D**), illustrating qualitatively similar kinetics and intracellular Ca^2+^ homeostatic pathways to those observed in supporting cells.

**Figure 4.**
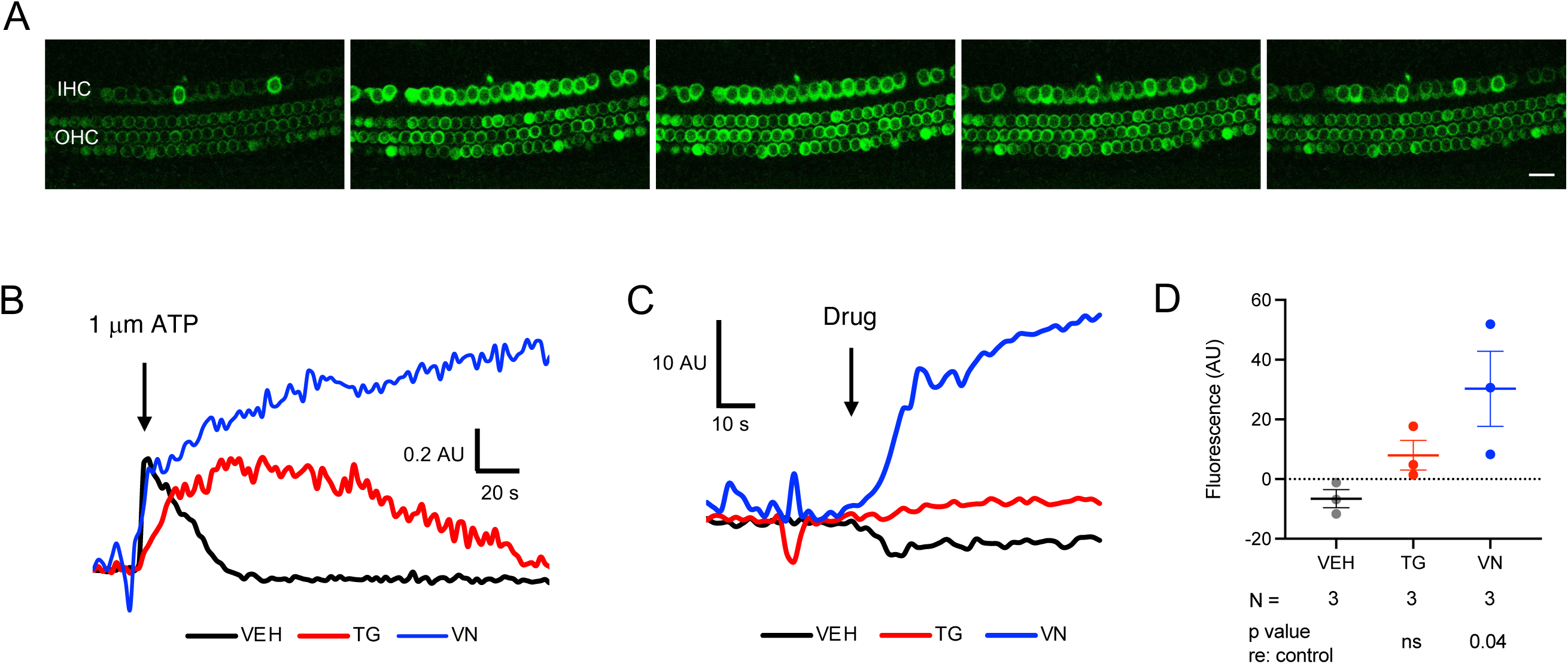
Ca^2+^ activity in neonatal cochlear hair cells. **A. Live imaging of in Myo15Cre-GcAMP neonatal cochlea.** No spontaneous Ca^2+^ peak activity is seen in hair cells of the neonatal cochlea. However, application of 1 μm ATP induced an increase in cytosolic Ca^2+^ with subsequent return to baseline in both inner hair cells (IHCs) and outer hair cells (OHCs). Time interval between successive images is 8s. Scale bar: 20 μm. **B. ATP-induced Ca^2+^ peak.** In an individual cochlear IHC, ATP treatment induced an increase in Ca^2+^ in the presence of vehicle (VEH, black), thapsigargin (TG, red), and vanadate (VN, blue), with prolonged onset and decay time with TG and prolonged elevation with no return to baseline in the presence of VN. Peak height is normalized to maximum amplitude of the initial ATP-induced peak. **C-D. Change in Steady-state Ca^2+^ after drug treatment.** Treatment with VN (blue) induced an in increase in Ca^2+^ across all IHCs and OHCs, as seen in mean fluorescence tracings across IHC and OHC regions **(C)** as well as mean fluorescence level change between baseline and drug treatment **(D).** Vehicle (VEH, black) treatment had no effect; Thapsagargin (TG) induced a slight increase in Ca^2+^. Means ± sem, with individual values in gray. N = 3 cochleae per condition. P values indicate pairwise comparisons relative to control (VEH) on 2-tailed unpaired Student’s t test. ns, not significant. AU, arbitrary units

### Live Ca^2+^ imaging in hair cells and supporting cells in the mature, hearing cochlea

Having established the ability to perform hair-cell and supporting-cell-specific live cytosolic Ca^2+^ imaging in neonatal cochlea, we moved to our juvenile (7-8-week-old), mature-hearing cochlear preparation, which enables imaging of the 8-10 kHz region of the cochlea (27,28). GcAMP-expressing mice have similar baseline hearing thresholds and response to 98-dB and 106-dB SPL noise exposures as wild-type CBA/J mice (**Supplemental Figure S2**). GcAMP expression was visible in both hair cells (in Myo15Cre-GcAMP mice) and supporting cells (in Sox2Cre-GcAMP mice), and cells remained stable in shape and size, with no steady-state changes in cytosolic Ca^2+^ levels over the recording period, suggestive of overall cellular health in this explant preparation (**Supplemental Videos 3-4**). There were no spontaneous intracellular Ca^2+^ transients seen in hair cells (**Supplemental Video 3**). In supporting cells, which exhibited robust spontaneous activity in both IS and OS regions of the neonatal cochlea, reduced spontaneous activity was observed in the mature-hearing cochlea (**Supplemental Video 4, Figure 5**). This is consistent with prior report, which demonstrated quiescence of spontaneous ICS wave activity after the onset of hearing around postnatal day 14 (23). Overall, these findings demonstrate that the mature-hearing cochlear explant preparation from Myo15Cre-GcAMP and Sox2Cre-GcAMP is healthy and enables detection of cytosolic Ca^2+^, but that overall Ca^2+^ activity is quiescent under control, non-noise-exposed conditions.

**Figure 5.**
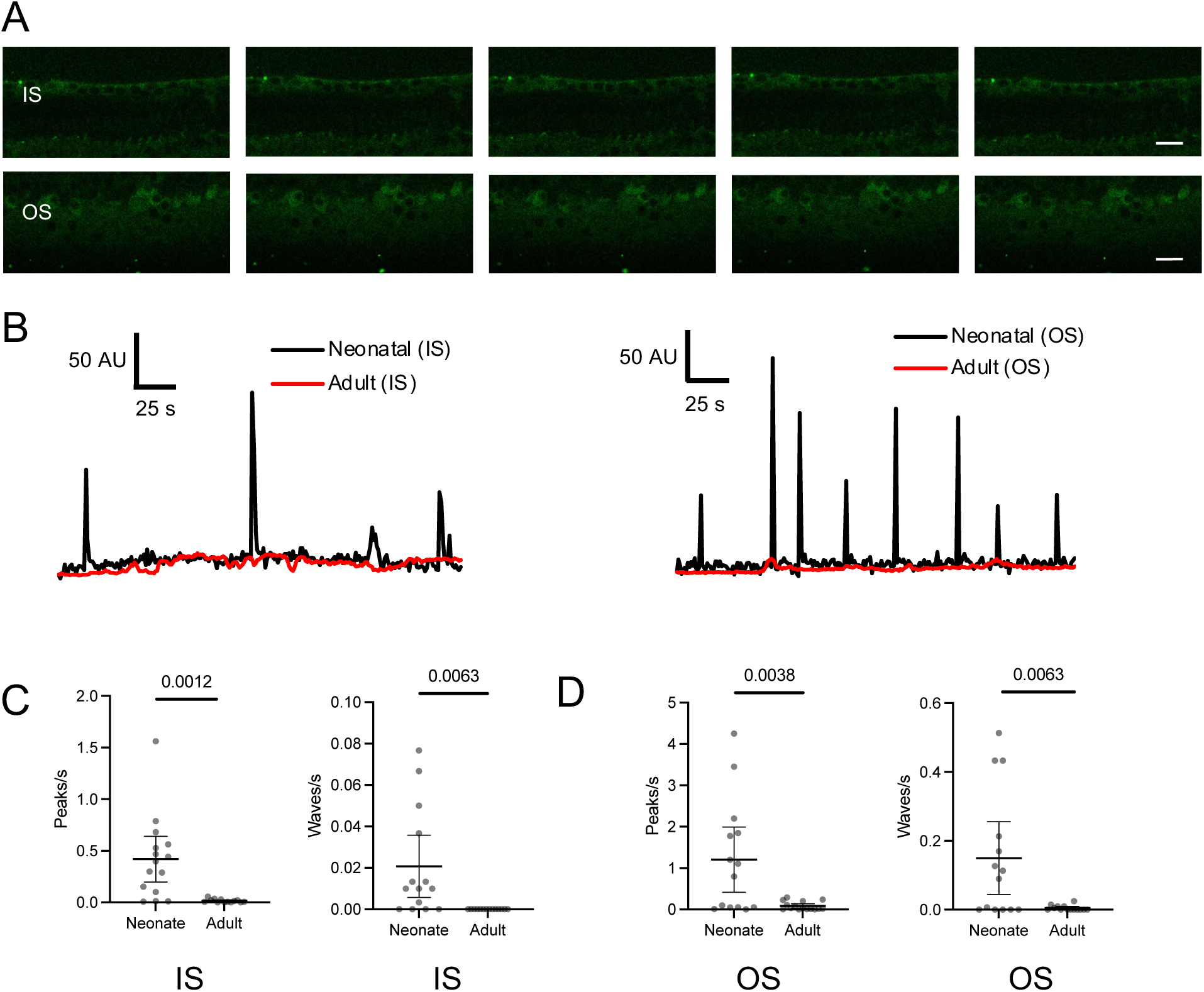
Ca^2+^ activity in adult cochlear supporting cells. **A. Live imaging of Sox2Cre-GcAMP adult cochlea.** Spontaneous Ca^2+^ peak activity is rarely seen in supporting cells of the adult cochlea. Time interval between successive images is 2s. Scale bar: 20 μm. **B-D. Activity in neonatal vs adult cochlea.** Compared with neonatal cochlea (black), single cells in the inner sulcus (IS, left) and outer sulcus (OS, right) of the adult cochlea (red) show no spontaneous Ca^2+^ peak activity. Number of peaks and waves per second were significantly higher in neonatal versus adult cochlea in both inner sulcus **(C)** and outer sulcus **(D)** regions. **C-D.** Means ± sem with individual values in gray. P values as indicated (black bars) on pairwise comparison between neonatal and adult cochlea using 2-tailed, unpaired Student’s t test. N = 15 (neonate) and 16 (adult) cochleae. AU, arbitrary units.

### Noise exposure activates ICS activity in cochlear supporting cells

Though spontaneous ICS waves in supporting cells of the mature-hearing cochlea were scant, noise exposure elicited abundant ICS activity (**Supplemental Video 5, Figure 6A**). ICS waves propagated across and between all supporting-cell types: phalangeal, inner and outer pillar, Dieters, Hensen’s and Claudius cells. 8-16 kHz octave-band noise at 98 and 106 dB SPL, which elicits TTS with cochlear synaptopathy and PTS with hair-cell death, respectively, both induced a significant increase in the number of cell-level Ca^2+^ transients as well as organ-level ICS waves within 1h of initiation of noise (**Figure 6B-C**). 24h after completion of a full 2h noise exposure, cochleae exposed to 106 dB SPL noise, which ultimately causes hair-cell death, had persistent Ca^2+^ peak activity, whereas those exposed to 98 dB SPL noise did not. For both noise exposure levels, the decay time of the Ca^2+^ transients in these supporting cells, which is a measure of persistent cytosolic Ca^2+^ elevation, was not immediately elevated, but became elevated 24h after noise exposure (**Figure 6D**). At the ICS wave level, cochleae exposed to 106 dB SPL noise had significantly longer distance of ICS wave propagation at both timepoints compared to both control and 98-dB SPL-exposed cochleae, which were not different from each other (**Figure 6E**). Taken together, these findings demonstrate that noise exposure induces ICS wave activity in cochlear supporting cells, and that this ICS-related cytosolic Ca^2+^ elevation is more persistent and extensive in cochleae exposed to the louder 106 dB SPL noise dose.

**Figure 6.**
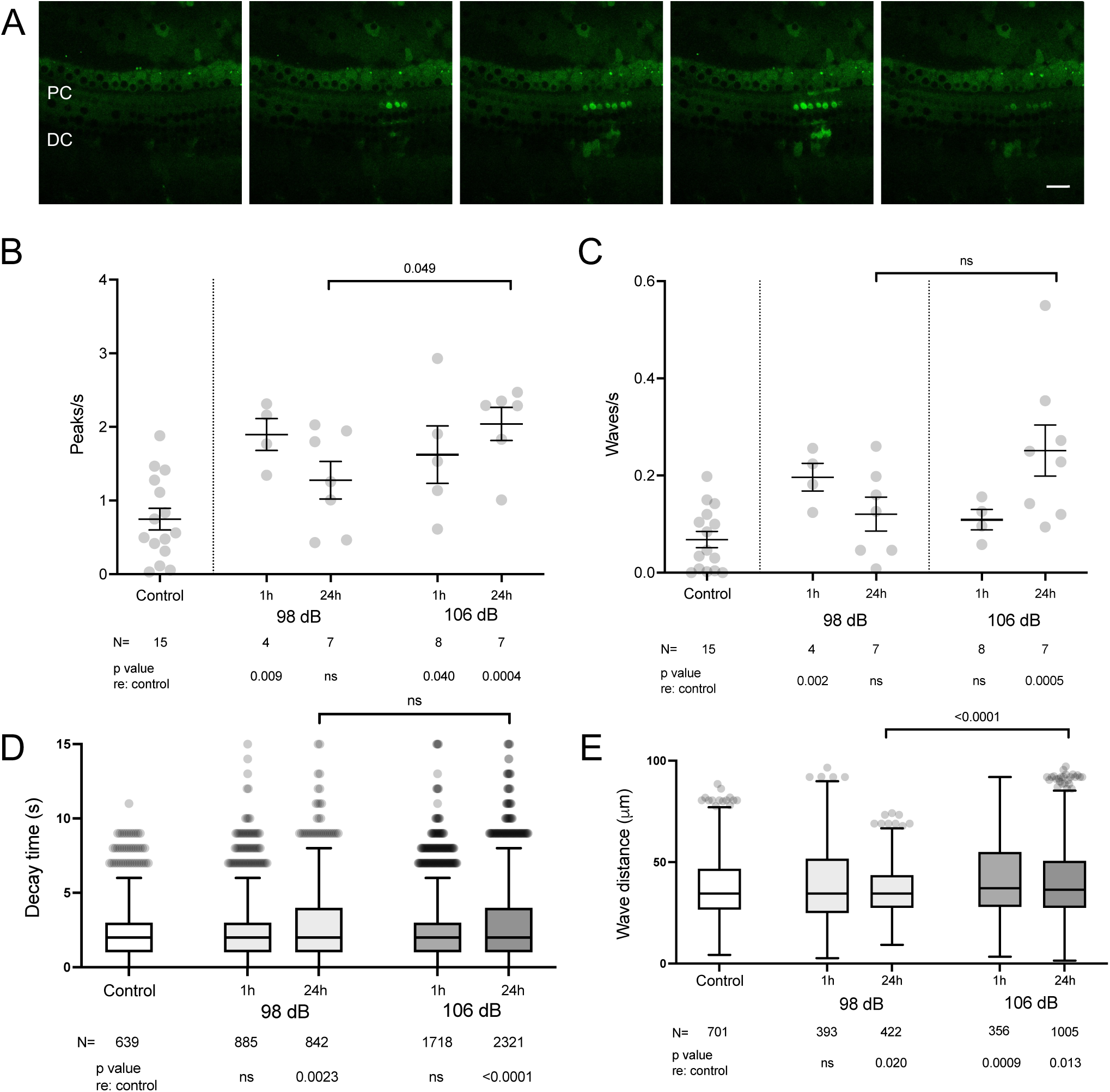
Ca^2+^ activity in adult cochlear supporting cells. **A. Live imaging of noise-exposed Sox2Cre-GcAMP adult cochlea.** Pre-exposure to 98-dB SPL noise induces ICS wave activity in supporting cells of the adult cochlea, including pillar cells (PC) and Dieter’s cells (DC). Time interval between successive images is 8s. Scale bar: 20 μm. **B-C. Effect of noise exposure on Ca^2+^ peak and ICS wave activity.** Compared with control, non-noise-exposed cochleae, cochleae from mice exposed to 98 dB as well as 106 dB noise showed increased number of Ca^2+^ peaks **(B)** and ICS waves **(C)** 1h after beginning of noise exposure. 24h after completion of noise exposure, 98-dB-exposed mice had no significant increase in either Ca^2+^ peaks or ICS waves compared to control, whereas 106-dB-exposed mice had persistent elevation in both Ca^2+^ peak activity and ICS waves. **D. Effect of noise exposure on Ca^2+^ peak decay time.** Ca^2+^ peak decay time is shown for individual Ca^2+^ peaks under the indicated conditions, demonstrating prolongation of Ca^2+^ peaks 24h after noise exposure after both 98 and 106-dB exposure. **E. Effect of noise exposure on ICS wave propagation distance.** Distance traveled for ICS waves was increased after 106-dB, but not 98-dB noise exposure, immediately after noise exposure. 24h later, 106-dB exposed cochleae had persistent increase in wave distance, whereas 98-dB cochleae had decreased propagation distance. **B-C.** Means ± sem, with individual values in gray. Sample size refers to the number of individual cochleae**. D-E.** Tukey plots are shown. Sample size refers to the number of individual peaks **(D)** or waves **(E)** analyzed. P values indicate pairwise comparisons on 2-tailed unpaired Student’s t test versus control, or specific pairs (black bars). ns, not significant.

### Cochlear hair cells are not responsive to noise or ATP but demonstrate cytosolic Ca^2+^- associated cell death after noise exposure

Adult cochlear hair cells displayed no spontaneous Ca^2+^ transients (**Supplemental Video 3**). Unlike neonatal hair cells, adult hair cells were not even responsive to external ATP. Application of 1 μM ATP elicited a robust Ca^2+^ transient in inner-sulcus supporting cells with off-target expression of GcAMP, but not the adjacent hair cells (**Figure 7A-B**). 106 dB noise exposure elicited occasional Ca^2+^ transients in OHCs (**Supplemental Video 6**). These transients could be differentiated into two types. Some transients had rapid onset and decay back to baseline, and were not associated with changes in hair-cell morphology (**Figure 7C**), though the decay times (27.2 ± 6.8 s (mean ± s.e.m.)) were significantly longer than those seen for Ca^2+^ transients in supporting cells (2.52 ± 0.06 s for 106-dB noise exposure, **Figure 6D**, p < 0.0001 compared to hair cells). Other OHC Ca^2+^ transients occurred over an even longer timecourse, and preceded hair-cell expansion and death (**Figure 7D**).

**Figure 7.**
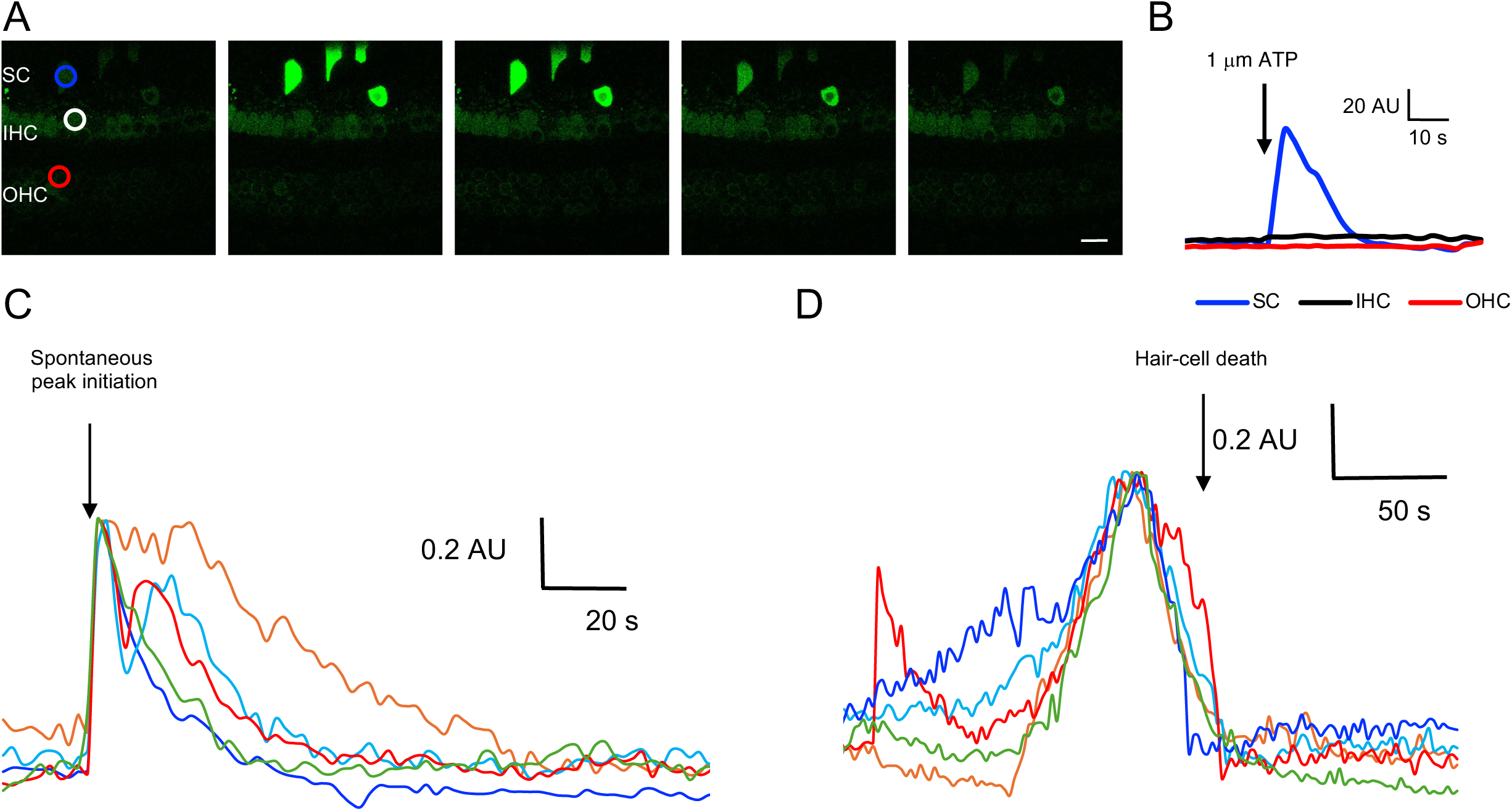
Live imaging of Myo15Cre-GcAMP adult cochlea. **A-B.** In the adult cochlea, no spontaneous Ca^2+^ peak activity is seen in hair cells **(A**). Though an inner-sulcus supporting cell (SC, blue) with off-target expression of GcAMP responded to ATP with a Ca^2+^ peak, neither an inner hair cell (IHC, white/black) nor an outer hair cell (OHC, red) was responsive. Time interval between successive images is 8s. Scale bar: 20 μm. **C-D. Noise-induced Ca^2+^ transients.** In fluorescence traces from OHCs from cochleae exposed to 106-dB noise, occasional cytosolic Ca^2+^ transients were observed. Transients with rapid rise and slow decay **(C)** were observed in OHCs that survived, whereas transients with slow rise and rapid decay **(D)** were observed in OHCs that subsequently expanded and died. Traces are normalized to maximal peak amplitude and aligned at the time of spontaneous peak initiation **(C)** or the time of hair-cell death **(D).** AU, arbitrary units.

## Discussion

In this study, we used a mature-hearing, physiologically relevant model of NIHL to evaluate critical components of ER stress – activation of the UPR and alterations of Ca^2+^ homeostasis within hair cells and supporting cells – in the cochlear response to acoustic overstimulation. We found that the UPR is indeed activated immediately after multiple levels of noise exposure, peaking within 2h (**Figure 2**), with a shift towards the pro-apoptotic PERK/CHOP pathway only with the 106 dB noise exposure level that causes permanent threshold shifts and hair-cell death (**Figure 1**). After demonstrating that a genetically-encoded Ca^2+^ indicator model that expresses GcAMP specifically in hair cells or supporting cells accurately reports cytosolic Ca^2+^ in neonatal cochleae (**Figures 3-4**), we studied the exact same noise-exposure models in mature-hearing, 7-8-week-old mice. Whereas neonatal, developing cochlea exhibited abundant spontaneous Ca^2+^ activity, especially ICS waves in supporting cells, both hair cells and supporting cells in the mature-hearing cochlea showed minimal spontaneous cytosolic Ca^2+^ transients or ICS waves, respectively (**Figure 5**). After noise exposure, however, supporting cells demonstrated increased ICS activity after noise exposure (**Figure 6).** 106 dB noise exposure, sufficient to cause permanent threshold shifts and pro-apoptotic UPR activation, was associated with more prolonged and extensive ICS wave activity in the 24h after noise exposure. In addition, some hair cells demonstrated an increase in cytosolic Ca^2+^ preceding hair-cell death (**Figure 7**). These transients were reminiscent of those observed in zebrafish hair cells exposed to aminoglycosides (13), suggesting a similar role for cytosolic Ca^2+^ accumulation in the events immediately preceding hair-cell death in the noise-exposed mammalian cochlea. Surviving hair cells generally did not demonstrate persistent elevation of cytosolic Ca^2+^, but instead showed only transient increases.

These findings — that noise exposure immediately induces ICS waves in supporting cells, Ca^2+^ transients in HCs, and UPR upregulation across the cochlea, with louder, PTS-associated noise specifically causing persistent ICS waves, UPR shift towards apoptosis, and cytosolic Ca^2+^ increases in HCs leading to their death— suggest that Ca^2+^ dysregulation and the UPR may constitute an early mechanism that can control subsequent hair-cell death and PTS. The effectiveness of ISRIB, a small-molecule eIF2B activator that specifically reduces the pro-apoptotic arm of the UPR, in preventing NIHL (6), further supports the notion that the UPR is causally involved in NIHL and can be targeted for treatment. This work demonstrates the need to understand more precisely the timeline and interrelationship of these cellular events and additional molecular mediators that might serve as targets for treatment. In particular, it remains unknown exactly how ICS waves in supporting cells interact with Ca^2+^ transients in hair cells. Do ICS waves in supporting cells induce hair-cell Ca^2+^ transients and, subsequently, cause hair-cell death? Or do dying hair cells induce ICS waves in the surrounding supporting cells? Indeed, the notion that ICS waves may be triggered by hair-cell damage is supported by studies on ICS waves in neonatal cultures (19,21), where mechanical trauma, laser ablation of hair cells, or neomycin treatment induced ICS waves and ERK1/2 activation in the supporting cells through which the ICS waves propagated. This ERK activation had further downstream effects on sensory epithelium remodeling and health of surrounding hair cells, illustrating the potential for ICS wave activity to modulate death and survival in the cochlea.

Alternatively, the opposite relationship is possible— ICS waves, which propagate in part through paracrine signaling mediated by ATP released by supporting cells (20), may trigger Ca^2+^ transients in adjacent hair cells. Whereas isolated transients may be tolerated by the hair cells, more intense or persistent ICS activity and ATP release may induce greater ER Ca^2+^ release in hair cells, thereby triggering hair cell death, either through ER Ca^2+^ depletion and the UPR or mitochondrial Ca^2+^ overload and oxidative stress. ATP is elevated in the endolymph after noise exposure (29), and targeting of purinergic receptor signaling has been proposed as a therapeutic strategy for NIHL (30). We found in this study that exogenous ATP induced robust cytosolic Ca^2+^ transients in neonatal hair cells but had no effect on hair cells in the mature-hearing cochlea. This may reflect changes in purinergic receptor over the course of hair-cell development (31,32); additionally, Ca^2+^ homeostasis is tightly regulated in hair cells between cytosolic buffering and highly regulated transfer between cytosol, ER, and mitochondria (12), and changes in cytosolic Ca^2+^ alone do not sufficiently predict cytotoxicity relating to downstream ER and mitochondrial effects (14). Further investigation of the precise interplay between ICS waves, ATP, and subcompartmental Ca^2+^ homeostasis in in mature, hearing cochlea is needed.

Nevertheless, the broad involvement of disorders of Ca^2+^ homeostasis, the UPR, and mitochondrial stress in genetic (12) and acquired (6,7,14) hearing loss highlight the need for further study to understand the underlying mechanisms in physiologically relevant disease models. In this study, we did not comprehensively evaluate all potential pathways by which noise exposure could induce changes in subcellular-compartment Ca^2+^ homeostasis and the associated downstream stress mechanisms. Specifically, we did not assess mitochondrial Ca^2+^ or stress pathways. However, our findings implicating cytosolic Ca^2+^ and the UPR in a consistent model of NIHL in mature-hearing mice does provide strong evidence for their involvement in the pathophysiology of NIHL. Understanding whether ICS waves are directly induced by noise and secondarily cause Ca^2+^ dysregulation in hair cells, or whether ICS waves are a response to hair cell injury that then subsequently helps to determine hair-cell fate, is critical in order to identify targets for treatment. More detailed investigation of cytosolic, mitochondrial, and ER Ca^2+^ homeostasis in cochlear hair cells, similar to that performed in zebrafish hair cells (13,14), as well as the downstream effects on mitochondrial and ER stress, in noise-exposed, mature-hearing mice is necessary to fully understand the roles of these pathways in hearing loss.

### Conclusion

In conclusion, we found that UPR activation and perturbations in cytosolic Ca^2+^ homeostasis in hair cells and supporting cells are involved in the cochlea’s early response to acoustic overstimulation. Given the critical role of the UPR, ICS waves, and cellular Ca^2+^ homeostasis in stress responses and subsequent cell fate, these findings suggest that targeting these pathways could be successful in treating NIHL. Further investigation into the specific mechanisms linking hair-cell and supporting-cell Ca^2+^ homeostasis and the UPR are necessary to more precisely identify targets for treatment.

## Methods

### Sex as a biological variable

Our study examined male and female animals, and similar findings are reported for both sexes.

### Mouse models and cochlear preparation

Sox2Cre (Jackson Laboratory #008454) or Myo15Cre (15) mice were bred with Ai95D mice (Jackson Laboratory #028865) for expression of GcAMP in supporting or hair cells, respectively. For wild-type noise exposures, 7-8-week-old CBA/CaJ mice (Jackson Laboratory, #000654) were used. Postnatal day 3-5 neonatal cochlear explant cultures were established as described (6) and used for live imaging. For live imaging in juvenile, mature-hearing cochleae (MS24), 7-8-week old mice were euthanized with carbon dioxide and decapitated immediately after noise exposure. The temporal bone was extracted from the skull by removal of the auditory bulla and then placed in ice-cold HBSS. Soft tissue and ossicles were removed with fine forceps and the otic capsule mounted on a custom 3D printed slide fixated in a hole on a plastic coverslip. Fresh HBSS was applied and the bone covering the apical turn of the cochlea was removed, preserving the membranous labyrinth and exposing the helicotrema. Reissner’s membrane was removed and the preparation used immediately for imaging. Time from euthanasia to imaging averaged less than 10 min.

Resonance (for Sox2Cre-GcAMP neonate) or line-scanning (for all other models) confocal imaging was performed on an upright Nikon A1R confocal microscope using a 60x water-immersion objective (NIR Apo, 1.0 NA), with temperature and CO_2_ control using a stagetop incubator (OKOlab). Optical sections in the x-y plane were recorded at 1x averaging, 1.2 AU pinhole, 1.1 dwell time, displaying the entire inner sulcus (IS), outer sulcus (OS) and hair cell regions in the middle cochlear turn (neonates) and apical cochlear turn (8-10 kHz region, juveniles).

For neonatal cultures, baseline imaging was performed for 3 minutes, followed by vehicle (media), thapsigargin (Tocris) and/or sodium orthovanadate (Calbiochem) application at t = 3 min and, for Myo15Cre-GcAMP only, ATP (Sigma) at t = 5 min. A total of 20 min imaging was performed, with interval between each successive image of 1 s for Sox2Cre-GcAMP and 2 s for Myo15Cre-GcAMP. For imaging of juvenile, mature-hearing, a single 10-min continuous imaging session was performed, with an interval of 2 s.

### UPR marker gene quantification

UPR marker gene expression was quantified in cochleae from noise-exposed animals as described (6). Briefly, at the indicated times after completion of noise exposure, animals were euthanized and cochleae harvested onto dry ice. Expression levels of three UPR markers (BiP, indicative of UPR activation; CHOP, correlated with pro-apoptotic activity of the UPR; and S-XBP1, associated with pro-homeostatic activity of the UPR) as well as GAPDH (as reference) were measured by qPCR and quantified against GAPDH and unexposed controls using the 2^-ΔCT^ method. Samples with inadequate GAPDH levels (defined as CT>24) were excluded from analysis. The ratio of CHOP over S-XBP1 was used as a marker of the pro-apoptotic state of the UPR (15).

### Image processing

Ca^2+^ fluorescence measurements were performed on regions of interest (ROIs) slightly larger than a cell, as done previously (6). Briefly, images were reoriented such that the hair cells were parallel to the bottom of the image. 325 ROIs, each 56 x 56 pixels (11 μm x 11 μm) in area were defined (ImageJ), and mean fluorescence intensity measured for every ROI at each timepoint. Ca^2+^ peak activity from each ROI-specific fluorescence timecourse was captured using a custom in-house script (Matlab, R2023a). Threshold for detection of a Ca^2+^ peak was set at 10 times the standard deviation of the baseline fluorescence measurement.

### Noise exposure and auditory testing

For NIHL induction, mice were exposed to 94, 98, or 106 dB SPL 8-16 kHz octave-band noise in a custom-built reverberant chamber as described (6), which respectively cause no hearing loss (94 dB), TTS with cochlear synaptopathy (98 dB) (16), and PTS with hair-cell death (106 dB) (6). Hearing was tested in mice by measuring auditory brainstem response (ABR) thresholds in response to broadband tone pips at 8, 16, and 32 kHz in the sound field using a standard commercial system (RZ6, Tucker-Davis Technologies) in a soundproof chamber as described (6).

### Experimental rigor and statistical analysis

Prior to analysis, outliers were removed in each dataset (ROUT method, Q=1%). For comparison between treatment groups for UPR gene expression, we used one-way ANOVA followed by post-hoc Dunnett’s multiple comparison tests. For pairwise comparison of Ca^2+^ dynamic parameters between groups, we used unpaired two-tailed Student’s t-test. Unless otherwise mentioned, results are presented as means +/- sem with sample sizes and p values between designated comparison groups as indicated in the figure legends, with a p-value <0.05 as significant, and lower p values indicated for specific comparisons. Statistical analyses were performed with GraphPad Prism 9.5.1. Sex was evaluated as a biological variable. UPR gene expression (**Figure 1**) and ABR thresholds (**Figure S1**) after different noise exposure levels were tested for male and female mice, and no significant differences found (unpaired two-tailed t-test with adjustment for multiple comparisons (Bonferroni correction for 3 simultaneous comparisons, for 3 noise-exposure levels). Because no differences were seen in these core measures between sexes, all subsequent analyses used pooled male and female mice.

## Study approval

This study was approved by the Institutional Animal Care and Use Committee of the University of California, San Francisco (AN1999783-00).

## Data Availability

All underlying data and supporting analytic code used in this study will be shared upon reasonable request.

## Author Contributions

Initial design: YP, JL, EHS, DKC. Experimental and ethical oversight, and funding: EHS and DKC. Experimental contributions: YP, JL, NIM, IRM, PS, and DKC. Data and statistical analysis: YP, JL, NIM, IRM, PS, and DKC. Manuscript drafting: YP, JL, and DKC. Manuscript review and approval: all authors. Multiple first and corresponding authors: YP and JL share first position, and EHS and DKC share final position. This work was a collaboration between groups focused on cell biology and genetics (conducted by JL and EHS) and auditory physiology and imaging (conducted by YP, NIM, IRM, PS and DKC). Because the physiology work constituted >50% of the actual content in the study, YP is listed first among the shared first position, and DKC is listed last among the shared final position.

## Acknowledgements

This study was funded by NIDCD R01 DC018583 (to DKC and EHS); and “Generation of cell lines to understand TMTC4, a novel human deafness gene,” from Hearing Research, Inc. (to DKC).

## References

1. Carroll YI, Eichwald J, Scinicariello F et al. Vital Signs: Noise-Induced Hearing Loss Among Adults in the United States 2011-2012. MMWR Morb Mortal Wkly Rep 2017; 66:139–144.

2. Yamane H, Nakai Y, Takayama M, Iguchi H, Nakagawa T, Kojima A. Appearance of free radicals in the guinea pig inner ear after noise-induced acoustic trauma. Eur Arch Otorhinolaryngol 1995; 252:504–508.

3. Kurabi A, Keithley EM, Housley GD, Ryan AF, Wong AC. Cellular mechanisms of noise-induced hearing loss. Hear Res 2017; 349:129–137.

4. Maeda Y, Fukushima K, Omichi R, Kariya S, Nishizaki K. Time courses of changes in phospho-and total-MAP kinases in the cochlea after intense noise exposure. PloS One 2013; 8(3):e58775.

5. Li P, Li S, Wang L, Li H, Wang Y, Liu H, Wang X, Zhu X, Liu Z, Ye F, Zhang Y. Mitochondrial dysfunction in hearing loss: Oxidative stress, autophagy and NLRP3 inflammasome. Front Cell Dev Biol 2023; 11:1119773.

6. Li J, Akil O, Rouse SL, McLaughlin CW, Matthews IR, Chan DK, and Sherr EH. Deletion of Tmtc4 activates the unfolded protein response and causes postnatal hearing loss. J Clin Invest 2018; 128(11):5150–5162.

7. Wang X, Zhu Y, Long H, Pan S, Xiong H, Fang Q, Hill K, Lai R, Yuan H, Sha SH. Mitochondrial Calcium Transporters Mediate Sensitivity to Noise-Induced Losses of Hair Cells and Cochlear Synapses. Front Mol Neurosci 2019; 11:469.

8. Wang J, Van De Water TR, Bonny C, de Ribaupierre F, Puel JL, Zine A. A peptide inhibitor of c-Jun N-terminal kinase protects against both aminoglycoside and acoustic trauma-induced auditory hair cell death and hearing loss. J Neurosci 2003; 23(24):8596–607.

9. Kopke R, Slade MD, Jackson R, Hammill T, Fausti S, Lonsbury-Martin B, Sanderson A, Dreisbach L, Rabinowitz P, Torre P 3rd, Balough B. Efficacy and safety of N-acetylcysteine in prevention of noise induced hearing loss: a randomized clinical trial. Hear Res 2015; 323:40–50.

10. Eshraghi AA, Aranke M, Salvi R, Ding D, Coleman JKM Jr, Ocak E, Mittal R, Meyer T. Preclinical and clinical otoprotective applications of cell-penetrating peptide D-JNKI-1 (AM-111). Hear Res 2018; 368:86–91.

11. Chang PH, Liu CW, Hung SH, Kang YN. Effect of N-acetyl-cysteine in prevention of noise-induced hearing loss: a systematic review and meta-analysis of randomized controlled trials. Arch Med Sci 2021; 18(6):1535–1541.

12. Richard EM, Maurice T, Delprat B. Calcium signaling and genetic rare diseases: An auditory perspective. Cell Calcium 2023; 110:102702.

13. Esterberg R, Hailey DW, Coffin AB, Raible DW, and Rubel EW. Disruption of intracellular calcium regulation is integral to aminoglycoside-induced hair cell death. J Neurosci 2013; 33(17): 7513–25.

14. Esterberg R, Hailey DW, Rubel EW, Raible DW. ER-mitochondrial calcium flow underlies vulnerability of mechanosensory hair cells to damage. J Neurosci 2014; 34(29):9703–19.

15. Li J, Choi BY, Eltawil Y, Ismail Mohamad N, Park Y, Matthews IR, Han JH, Kim BJ, Sherr EH, Chan DK. TMTC4 is a hair cell-specific human deafness gene. JCI Insight 2023; 8(24):e172665

16. Rouse SL, Matthews IR, Li J, Sherr EH, Chan DK. Integrated stress response inhibition provides sex-dependent protection against noise-induced cochlear synaptopathy. Sci Rep 2020; 10(1):18063.

17. Scemes E, Giaume C. Astrocyte calcium waves: what they are and what they do. Glia 2006; 54(7):716–725.

18. Leybaert L, Sanderson MJ. Intercellular Ca(2+) waves: mechanisms and function. Physiol Rev 2012; 92(3):1359–92.

19. Gale JE, Piazza V, Ciobotaru CD, Mammano F. A mechanism for sensing noise damage in the inner ear. Curr Biol 2004; 14(6):526–9.

20. Majumder P, Crispino G, Rodriguez L, Ciubotaru CD, Anselmi F, Piazza V, Bortolozzi M, Mammano F. ATP-mediated cell-cell signaling in the organ of Corti: the role of connexin channels. Purinergic Signal 2010; 6(2):167–87.

21. Lahne M and Gale JE. Damage-induced activation of ERK1/2 in cochlear supporting cells is a hair cell death-promoting signal that depends on extracellular ATP and calcium. J Neurosci 2008; 28(19):4918–28.

22. Tritsch NX, Yi E, Gale JE, Glowatzki E, Bergles DE. The origin of spontaneous activity in the developing auditory system. Nature 2007; 450(7166):50-5.

23. Tritsch NX and Bergles DE. Developmental regulation of spontaneous activity in the mammalian cochlea. J Neurosci 2010; 30(4):1539–1550.

24. Sirko P, Gale JE, Ashmore JF. Intercellular Ca^2+^ signalling in the adult mouse cochlea. J Physiol 2019. 597(1):303–317.

25. Piazza V, Ciubotaru CD, Gale JE, Mammano F. Purinergic signalling and intercellular Ca^2+^ wave propagation in the organ of Corti. Cell Calcium 2007; 41(1):77–86.

26. Chan DK, Rouse SL. Sound-Induced Intracellular Ca^2+^ Dynamics in the Adult Hearing Cochlea. PLoS One 2016; 11(12):e0167850.

27. Ismail Mohamad N, Santra P, Park Y, Matthews IR, Taketa E, Chan DK. Synaptic ribbon dynamics after noise exposure in the hearing cochlea. Accepted for publication, Comms Bio.

28. Müller M, von Hünerbein K, Hoidis S, Smolders JW. A physiological place-frequency map of the cochlea in the CBA/J mouse. Hear Res 2005; 202(1-2):63–73.

29. Muñoz DJ, Kendrick IS, Rassam M, Thorne PR. Vesicular storage of adenosine triphosphate in the guinea-pig cochlear lateral wall and concentrations of ATP in the endolymph during sound exposure and hypoxia. Acta Otolaryngol 2001; 121(1):10–5.

30. Berekméri E, Szepesy J, Köles L, Zelles T. Purinergic signaling in the organ of Corti: Potential therapeutic targets of sensorineural hearing losses. Brain Res Bull 2019; 151:109–118.

31. Jovanovic S, Milenkovic I. Purinergic Modulation of Activity in the Developing Auditory Pathway. Neurosci Bull 2020; 36(11):1285–1298.

32. Babola TA, Li S, Wang Z, Kersbergen CJ, Elgoyhen AB, Coate TM, Bergles DE. Purinergic Signaling Controls Spontaneous Activity in the Auditory System throughout Early Development. J Neurosci 2021; 41(4):594–612.

