## Supplementary figures and images for "Noise induces intercellular Ca^2+^ signaling waves and the unfolded protein response in the hearing cochlea"

### Supplemental Figure 1

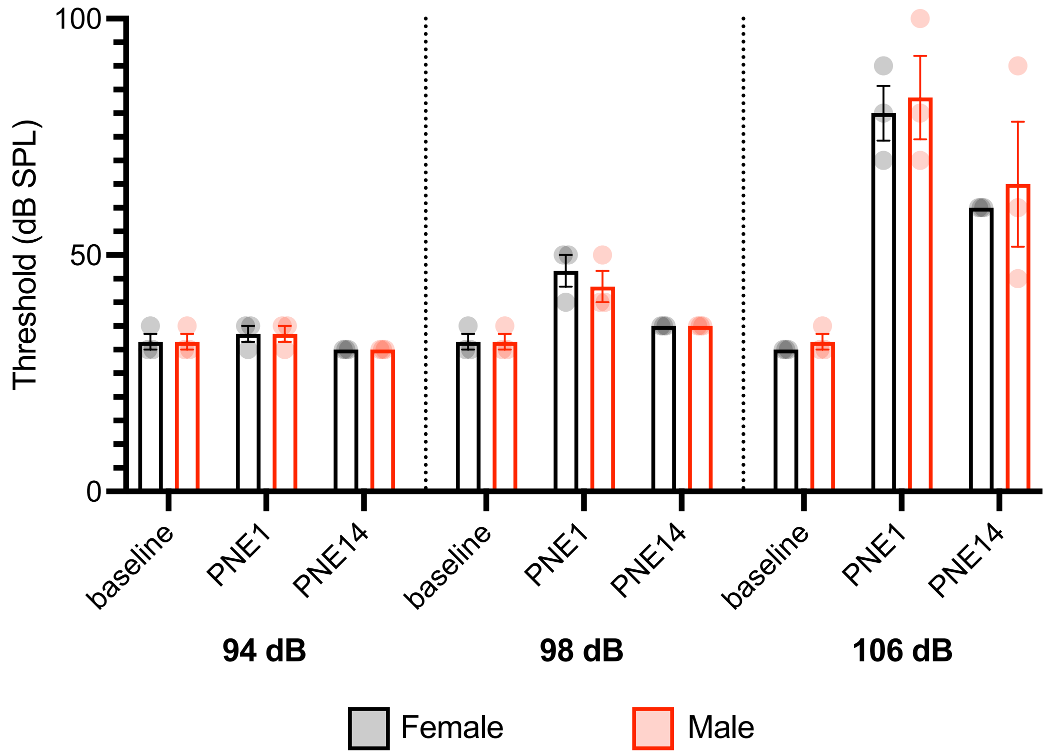

### Supplemental Figure 2

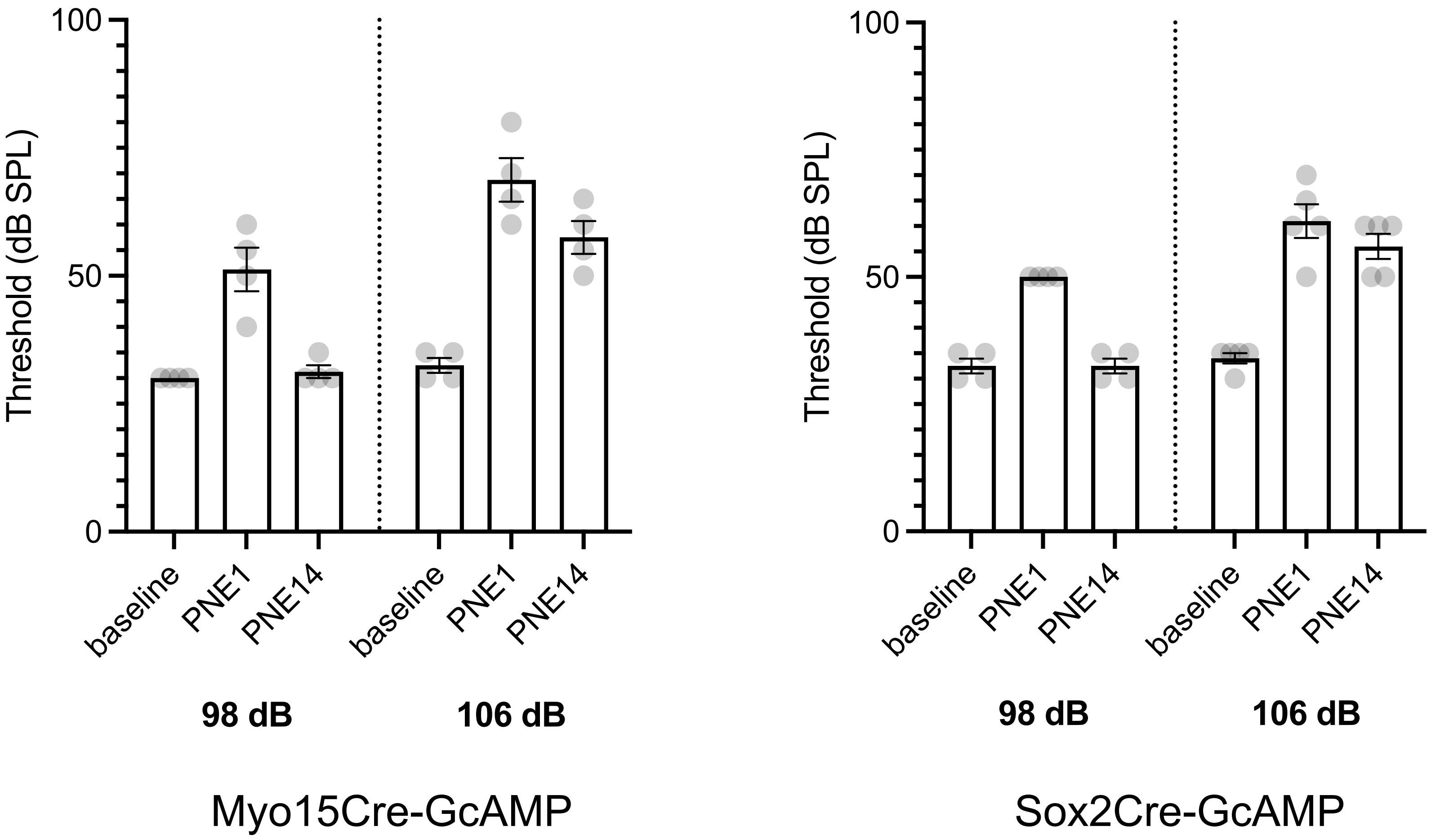
